# TimeTrial: An Interactive Application for Optimizing the Design and Analysis of Transcriptomic Times-Series Data in Circadian Biology Research

**DOI:** 10.1101/2020.04.15.043695

**Authors:** Elan Ness-Cohn, Marta Iwanaszko, William Kath, Ravi Allada, Rosemary Braun

## Abstract

The circadian rhythm drives the oscillatory expression of thousands of genes across all tissues, coordinating physiological processes. The effect of this rhythm on health has generated increasing interest in discovering genes under circadian control by searching for periodic patterns in transcriptomic time-series experiments. While algorithms for detecting cycling transcripts have advanced, there remains little guidance quantifying the effect of experimental design and analysis choices on cycling detection accuracy. We present TimeTrial, a user-friendly benchmarking framework using both real and synthetic data to investigate cycle detection algorithms’ performance and improve circadian experimental design. Results show that the optimal choice of analysis method depends on the sampling scheme, noise level, and shape of the waveform of interest; and provides guidance on the impact of sampling frequency and duration on cycling detection accuracy. The TimeTrial software is freely available for download and may also be accessed through a web interface. By supplying a tool to vary and optimize experimental design considerations, TimeTrial will enhance circadian transcriptomics studies.

## Introduction

The circadian rhythm, observed as a periodic 24 hour behavioral and physiological cycle at the organismal level, is governed by an evolutionarily–conserved set of core clock genes operating at the transcriptional and protein level. Consisting of only a few genes, the circadian clock coordinates a vast array of cellular processes, including the cyclic expression of nearly half of genes across all tissues (Zhang et al., 2014). This rhythm can be entrained to environmental cues (Zeitgebers) such as light, temperature, and food, allowing external stimuli to modulate time–of–day specific functions. While numerous epidemiological studies have demonstrated significant links between circadian rhythms and human health (Braun et al., 2018; Chang et al., 2009; Kathale and Liu, 2014; Levi and Schibler, 2007; Patke et al., 2017; Puttonen et al., 2010; Roenneberg et al., 2007; Videnovic et al., 2014; Zhang et al., 2014), the underlying mechanisms linking the circadian clock to health outcomes remain largely unknown.

The advent of high-throughput omic technology now enables researchers to investigate these mechanisms in molecular detail by tracking the expression of thousands of transcripts over the course of the day, with the goal of identifying specific genes under circadian control. However, there are a number of analytical challenges in extracting rhythmic signals from noisy transcriptomic data. First, experimental limitations constrain the frequency and length of sampling, requiring inferences to be made from sparse or short time-series measurements. Second, cycling genes’ expression profiles often do not exhibit sinusoidal trajectories; sharp peaks, damped oscillations, and additive linear trends have all been observed.

To address the complexities of circadian rhythm detection, a range of non-parametric methods have been developed (Hughes et al., 2010; Hutchison et al., 2018; Perea et al., 2015; Thaben and Westermark, 2014; Wu et al., 2016; Yang and Su, 2010). These methods search for evidence of periodicity (e.g., by testing for correlations with template waveforms (Hughes et al., 2010; Hutchison et al., 2018)). However, subsequent benchmarking studies demonstrate that different methods can yield conflicting results when run on the same dataset (Deckard et al., 2013; Serpedin et al., 2008; Wu et al., 2014). Moreover, performance depends on the shape of the signal being detected, noise levels, and sampling schemes (Deckard et al., 2013; Hughes et al., 2009).

In addition to performance differences, there are also considerations of various methods’ abilities to handle replicates, uneven sampling, missing data, and computational efficiency. In practice, the ability for methods to adequately handle these features directly impacts researcher’s flexibility in experimental design. For instance, a method that can accommodate uneven sampling can allow for dense sampling at times of interest, with sparser sampling at other times. Because missing data often occurs as a result of sequencing errors with greater likelihood as sample size increases (Gierliński et al., 2015), researchers benefit from algorithms that can handle missingness without the need to impute data. Finally, computational efficiency allows for dataset sizes to grow, while still processing the data in a reasonable amount of time. Taken together, methodological constraints imply that the choice of cycling detection method will necessarily impact the optimal experimental design, and vice versa.

These considerations, coupled with the need to limit costs, imply that designing an optimal circadian time–series experiment is a non-trivial task. While recommendations for experimental designs have been made (Hughes et al., 2007, 2017; Wu et al., 2014), quantitative tools to to flexibly and comprehensively weigh these considerations in the context of real data remain lacking. Moreover, while researchers have attempted to define criteria for method usage (Deckard et al., 2013; Hutchison et al., 2018; Wu et al., 2014), no guidance exists for custom sampling schemes, as previous studies used fixed sampling schemes and a limited number of waveform shapes.

To address these challenges, this article focuses on optimizing circadian rhythm detection by introducing a framework to evaluate the reliability of cycling detection as sampling schemes, waveform shapes, and cycling detection algorithms are varied. The results provide valuable evidence–based guidance for experimental design and analysis choices. As part of this work, we developed TimeTrial: an interactive, user–friendly, open– source software suite that enables circadian researchers to perform head-to-head comparisons of four leading cycle detection methods (JTK_Cycle (Hughes et al., 2010), ARSER (Yang and Su, 2010), RAIN (Thaben and Westermark, 2014), and BooteJTK (Hutchison et al., 2018); **Supplementary Table 1**) using both synthetic and real data. With TimeTrial, researchers can further explore these methods’ performance under different noise levels, number of replicates, length of sampling, sampling resolution, and waveform shapes. An innovative feature of TimeTrial is the ability for the researcher to specify an arbitrary custom sampling scheme and obtain comparison of the cycling detection results, allowing them to gauge how their design choices may impact the experimental findings. Together, these results will enhance rigor and reproducibility in future circadian time–series experiments and improve tools to analyze cycling genes in increasingly large and complex datasets.

## Results

TimeTrial was developed as a tool for the design and optimization of omic time-series experiments in circadian biology research. Consisting of two interactive applications using both synthetic and real data, TimeTrial allows researchers to explore the effects of experimental design on cycling detection, examine the reproducibility of cycling detection methods across biological datasets, and optimize experimental design for cycle detection. Applied to four cycle detection methods (JTK_Cycle (Hughes et al., 2010), ARSER (Yang and Su, 2010), RAIN (Thaben and Westermark, 2014), and BooteJTK (Hutchison et al., 2018); **Supplementary Table 1**), our results reveal that no method consistently outperforms all others in all circumstances, but rather that the performance depends on the sampling schemes and waveforms of interest. An interactive interface allows researchers to explore performance under different sampling schemes (including varying lengths, resolutions, and irregular sampling), providing valuable guidance for the optimization of circadian transcriptomic experiments given practical constraints (e.g., number of samples) and the signals of interest.

TimeTrial provides insights into cycling detection performance using both synthetic data and real data. The synthetic data provides precise control over the input data dynamics and noise, allowing the accuracy of cycling detection to be directly assessed when the ground truth (cycling/non-cycling) is known. The real data, which comes from multiple studies, allows cycling detection methods to be assessed in terms of the reproducibility of the findings in biologically representative datasets.

### Synthetic Data: Simulating Gene Expression Dynamics

To comprehensively evaluate the performance of cycling detection methods for different patterns of temporal gene expression, we systematically created synthetic datasets consisting of varying number of replicates, sampling intervals, sampling lengths, and noise levels for a variety of waveform shapes; in total, these represent 240 combinations of conditions (**Figure 1A**). 1000 “genes” were simulated with varying amplitudes, phases, and shape parameters (e.g., the envelope for damped/amplified waves) for each of the 11 base waveforms (**Figure 1B**), yielding in total 11,000 simulated genes for each of the 240 conditions. The choice of waveform shapes was inspired by patterns observed in experimental circadian datasets (**Supplementary Figure 1**), and is designed to give the user an avenue to explore the types of patterns that would be classified as cycling or non-cycling for various sampling and analysis choices. Further details of the synthetic datasets can be found in the *Methods* section and the *Supplement*.

**Figure 1:**
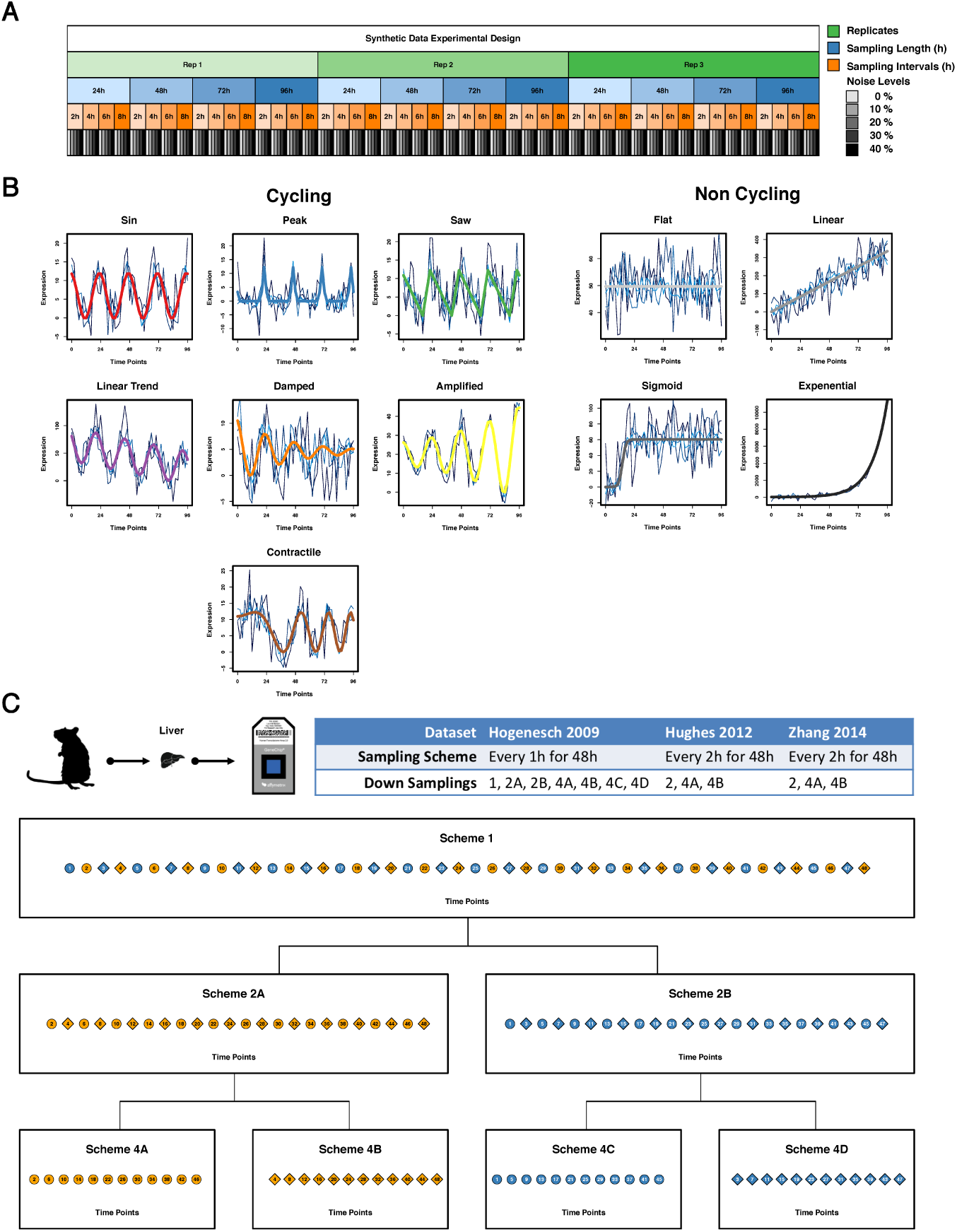
Experimental Design - Synthetic & Biological Data. [**A**] 240 unique synthetic time course datasets where generated in R to determine ground truth. Each dataset consisted of different number of replicates, sampling intervals, sampling lengths, and noise levels as a percent of the wave form amplitude. Each outer node represents an independent experiment. [**B**] Each dataset consisted of 11 different classes of waveforms commonly seen in real biological data. 7 were classified as cycling, and 4 as non-cycling. Signals are shown with additive gaussian noise as a percent of the wave form amplitude (i.e. 10%, 20%, 30%, & 40%). All cycling waveforms had a set period of 24 hours, except contractile which varied over time. [**C**] Three mouse liver time-series expression sets were analyzed from the Gene Expression Omnibus database: Hogenesch 2009 (GSE11923), Hughes 2012 (GSE30411), and Zhang 2014 (GSE54650). The Hughes and Zhang studies, sampled every 2h for 48h, were downsampled to 2 datasets sampled every 4 hour (Hughes_4A and Hughes_4B; Zhang_4A and Zhang_4B). The Hogenesch study, sampled every 1h for 48h, was downsampled to 2 datasets sampled every 2 hours (Hogenesch_2A and Hogenesch_2B) and also into 4 datasets sampled every 4 hours (Hogenesch_4A, Hogenesch_4B, Hogenesch_4C, and Hogenesch_4D).

### Biological Data: Reproducibility Analysis

While synthetic data has the benefit of having a known ground truth, it has the drawback of not necessarily being representative of real biological datasets. On the other hand, measuring a method’s accuracy using real data is limited as the ground truth is not generally known. Instead, however, one may test the *reproducibility* of the results, under the assumption that a true biological signal should be consistently detected across multiple studies of the same condition.

To this end, we took a “cross–study concordance” approach in which we tested methods’ ability to consistently characterize a set of 12868 genes measured in three independent studies as cycling or noncycling. By evaluating the rank correlation *ρ* of the cycling detection *p*-values obtained from the various studies, our analysis directly quantifies reproducibility. We analyzed three distinct mouse liver time–series expression sets (Hughes et al., 2009, 2012; Zhang et al., 2014) to evaluate the concordance. Additionally, we downsampled each dataset to mimic the effect of sparser sampling. Additional details can be found in the *methods* section.

### Experimental Design Recommendations

Using TimeTrial to analyze common sampling schemes in both synthetic and biological datasets, the following sections present recommendations for the experimental design framework in regards to selection of sampling scheme and concatenation. For the development of customized sampling schemes that may better fit the individual researcher’s needs, users are encouraged to explore TimeTrial’s custom sampling feature as described in the *TimeTrial Capabilities and Usage* section.

#### Sampling Resolution

While long, frequently–sampled timeseries provide the clearest picture of circadian dynamics, this must be balanced with practical considerations such as experimental cost. It is thus of interest to identify an optimal sampling scheme by varying sampling length, resolution, and replicates. To determine the limits of cycling detection as the frequency and length of sampling is reduced, we downsampled three datasets and compared the results across all genes (**Figure 1C**).

To determine whether a method gives consistent results at lower sampling, we compute the rank correlation *ρ* of the cycling detection *p*-values across all genes for different data sets and sampling schemes to quantitatively determine whether two methods yield the same *ranking* of cycling genes, without setting arbitrary *p*-value thresholds. Results reveal that 1-hour and 2-hour sampling resolutions show high correlation across all datasets and methods with *ρ* values between 0.79 − 0.94 (**Supplementary Figure 2**). This implies that at 1- and 2-hour sampling, all methods reproducibly rank the same genes (from most to least cycling) in the three independent datasets. It also suggests that different methods identify the same genes at the top of their respective lists, implying that the choice of methods does not significantly affect the results at these resolutions. In contrast, the results from 4-hour sampling resolutions show poorer correlations across datasets and methods, with *ρ* values between 0.67 − 0.89 (**Supplementary Figure 2**), implying that results become less reliably reproducible with 4-hour sampling. Moreover, when working in the low density regimes, there is less concordance between methods, suggesting that the choice of method has a large impact on the cycle detection results (**Supplementary Figure 3**). Taken together, these results suggest that cycle detection is robust at 1- and 2-hour sampling intervals but becomes significantly less reliable at 4-hour sampling intervals.

Importantly, we find that 4-hour sampling not only “misses” cycling genes that are detected at 1- and 2-hour sampling (false negatives); it can also lead to erroneously calling non-cycling genes as cycling (false positives). **Figure 2A** illustrates a gene that is not rhythmic in the 2-hour sampling but appears periodic when down–sampled to 4 hours. A cluster of such genes can be seen in the lower right of the scatter plots in (**Figure 2B**), where genes have a non-significant − log(*p*) in the 2-hour sampling scheme, but a significant − log(*p*) the 4-hour sampling scheme. **Figure 2C** shows the overlap of genes detected as cycling at (FDR<0.05) under 2– and 4–hour sampling. Additionally, we investigated whether known core clock genes were consistently detected as cycling; while they are robustly detected in the 1–hour and all 2–hour datasets, they are less reliably detected in the 4–hour data, further corroborating the findings above.

**Figure 2:**
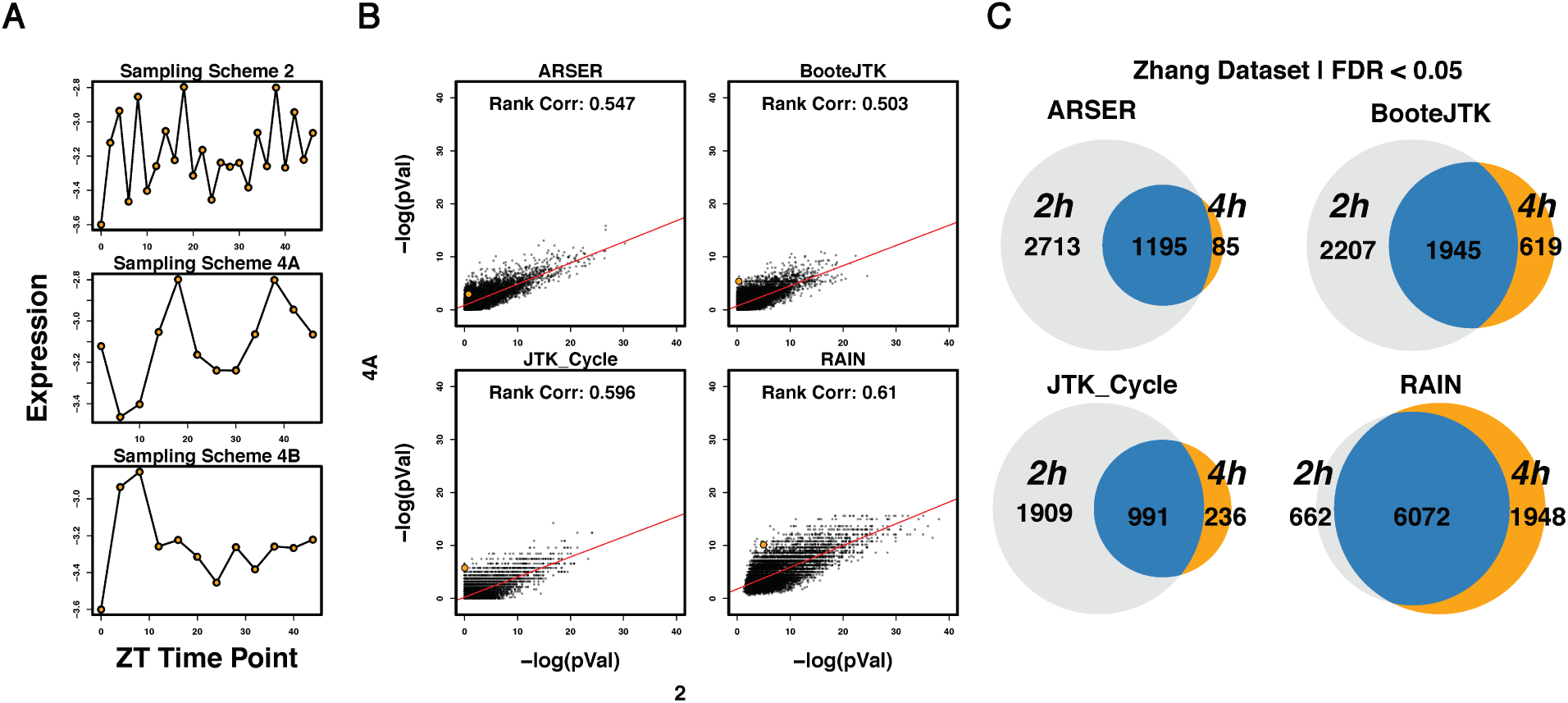
Experimental Resolution. [**A**] Expression time courses of P2ry10b gene at 2 hour and the two downsampled 4 hour (4A and 4B) resolution from the Zhang Dataset. [**B**] Scatter-plots of −log(pVals) of 2 hour sampling versus 4 hour sampling across all methods. p-values are not FDR corrected for comparison purposes. The orange point denotes the P2ry10b gene highlighted in the right panel. [**C**] Venn diagram of genes detected as cycling (FDR adjusted p-val < 0.05) in the 2 hour versus 4 hour down sampled Zhang dataset across all methods. Genes detected as cycling in the 4A and 4B conditions were grouped together.

One may then ask whether it is better to devote resources to more frequent sampling, or to a greater number of replicates at lower sampling rates. From the standpoint of sequencing cost, sampling every 4-hours for 48 hours with 2 replicates is the same as sampling every 2-hours for 48 hours with 1 replicate. Tests with synthetic data show these two schemes are similarly powered in their ability to detect true cycling genes; however, the 2-hour single-replicate scheme has the benefit of fewer false positives compared 4-hour duplicate sampling (**Figure 2A**). These findings suggest that sampling every 2 hours for 48 hours with a single replicate is advantageous over sampling every 4 hours in duplicate.

#### Concatenation Bias

Concatenation of replicate time-series is common practice in the field of circadian biology(Duong et al., 2011), whereing researchers will concatenate two replicate 24-hour series into a single 48-hour series. This is done under the assumption that if a signal has a true 24 hour period, concatenation of the signal will maintain its cyclic nature. However, concatenation of replicates can induce apparent 24-hour periodicity for nonperiodic signals (**Figure 3A**).

**Figure 3:**
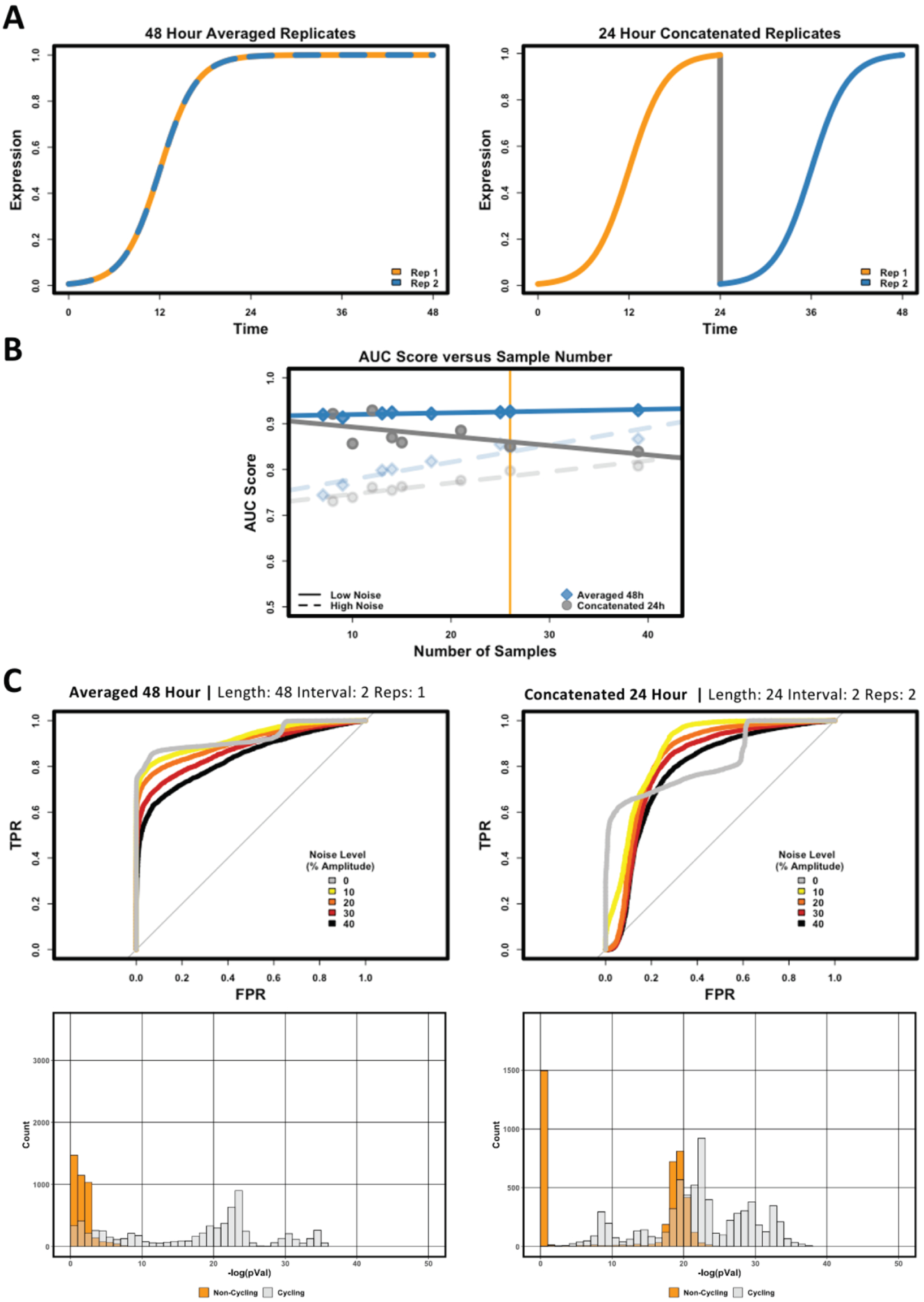
Concatenation Bias. [**A**] Cartoon representation of concatenated versus averaged replicates of sigmoidal waveform. **TOP:** Non-oscillatory when sampled for 48 hour with averaged replicates. **Bottom:** Artificially oscillatory when sampled for 24 hour with concatenated replicates. [**B**] AUC scores of the 24 hour and 48 hour time-courses from the synthetic datasets were plotted as a function of the number of samples using BooteJTK. Time courses varied in sampling interval and number of replicates. 24 hour time-courses were concatenated; while 48 hour time courses were averaged across replicates. Solid lines represent AUC scores in low noise regimes(0% and 10%) and dashed lines represent AUC scores in high noise regimes(30% and 40%). The vertical line represents the samples highlighted in panel B of the figure. [**C**] **TOP:** ROC curves for all noise levels at the 48 hour every 2 hour with 1 rep and 24 hour every 2 hour with 2 reps sampling scheme. Both schemes have the same number of timepoints. **BOTTOM:** Histograms of −log(pVals) of cycling (light grey) and non-cycling (orange) waveforms in each condition. Larger −log(pVals) denotes waveforms detected as more significantly cycling by BooteJTK.

As we increase sampling resolution in the low noise regime, accuracy for the concatenated data *decreases*, as seen by the negative slope in **Figure 3B** (solid grey line). Paradoxically, this means our ability to correctly classify genes becomes worse with more samples with less noise. A closer look at the samples highlighted by the orange line in **Figure 3B** illustrates the reason for this effect (*24h at 2h with 2 replicates* vs. *48h at 2h with 1 replicate*, **Figure 3C**). The ROC curves show that concatentation of two 24-hour replicates increases the false positive rate (FPR) over the 48 hour case due to periodic replication of any transient dynamics in the first 24 hour period (**Figure 3C**). As we increase our sampling resolution, we can better discern signal shape; coupled with concatenating expression patterns at the 24 hour mark, this increased clarity causes methods searching for periodicity with a period of 24 hours to erroneously classify sigmoidal, linear, and exponential signals as rhythmic. This finding corroborates other published studies (Hughes et al., 2017, 2009) that have likewise strongly recommended against concatenation. Taken together, these results imply time-series data should not be concatenated prior to statistical testing.

### TimeTrial Capabilities and Usage

The optimal choice of cycling detection method depends on sampling scheme, noise level, number of missing data-points, number of replicates, and shape of the waveform of interest. Thus, in addition to the above recommendations, we provide TimeTrial as a freely available tool for users to explore cycling detection performance and to optimize their sampling schemes.

As described above, TimeTrial consists of two components: one that analyzes cycling detection accuracy using synthetic data, and another that analyzes cycling detection reproducibility using real data. In the first TimeTrial component, using synthetic data, users can experiment with different number of replicates, sampling lengths, sampling resolutions, and noise levels to assess their effects on detection of specific signal shapes (**Figure 4**). By comparing the reported ROC curves, AUC scores, and *p*-value distributions for each different sampling scheme and methods, users can determine the optimal sampling scheme and method for cycle detection of various waveform (i.e. cosine, peak, saw-tooth, etc.). Furthermore, users can assess the robustness of each method when noise is introduced by comparing the standard deviation AUC scores. Given a specified sampling scheme, methods with lower standard deviations in AUCs imply the cycling detection results are stable across increasing noise values, while larger standard deviations indicate results are more strongly influenced by the noise level.

**Figure 4:**
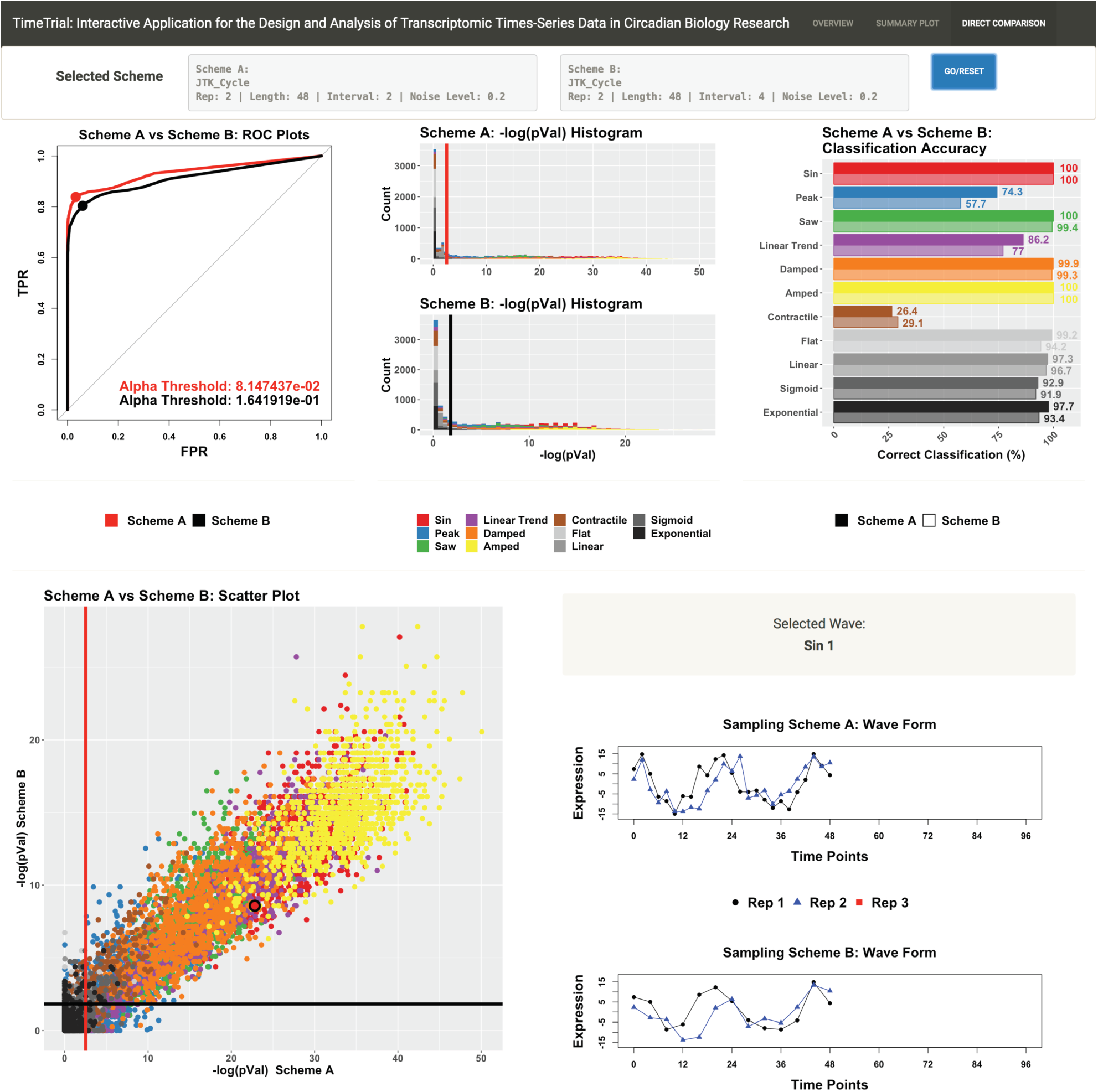
Synthetic Data - TimeTrial: Interactive Application for Circadian Rhythm Study Design. The synthetic dataset version of TimeTrial allows users to directly compare methods across different sampling lengths, sampling intervals, number of replicates, and noise levels. Users can further set different significance thresholds and take closer looks into the ability of methods to detect different waveform patterns. See (https://github.com/nesscoder/TimeTrial and/or https://nesscoder.shinyapps.io/TimeTrial_Synthetic/) for interactive plots and a complete tutorial.

Additionally, users can determine how their choices affect the *p*-value distribution of signal shapes to determine how robust a sampling scheme and method are at separating specific types of cycling signals from non-cyclers. Finally, users can adjust *p*-value thresholds and inspect output on a per signal basis to help determine appropriate threshold cutoffs for defining separation between signal shapes for downstream analysis.

In the second TimeTrial component, using real data, users can compare sets of genes detected as cycling by different methods and under different sampling schemes (**Figure 5**). By comparing results across each dataset and downsampling, users can perform a concordance analysis to determine which genes are picked up by each method and each sampling scheme. The TimeTrial interface allows the user to explore cycling detection results at various levels of significance and minimal fold–change (a criterion commonly used to reduce false positives). Given the knowledge of how sampling scheme effects the detection of waveform shape from use of the synthetic datasets, users can test the ability to pick up these shapes in the biological dataset and judge whether the waveform shapes being detected as cycling are representative of the patterns they wish to detect.

**Figure 5:**
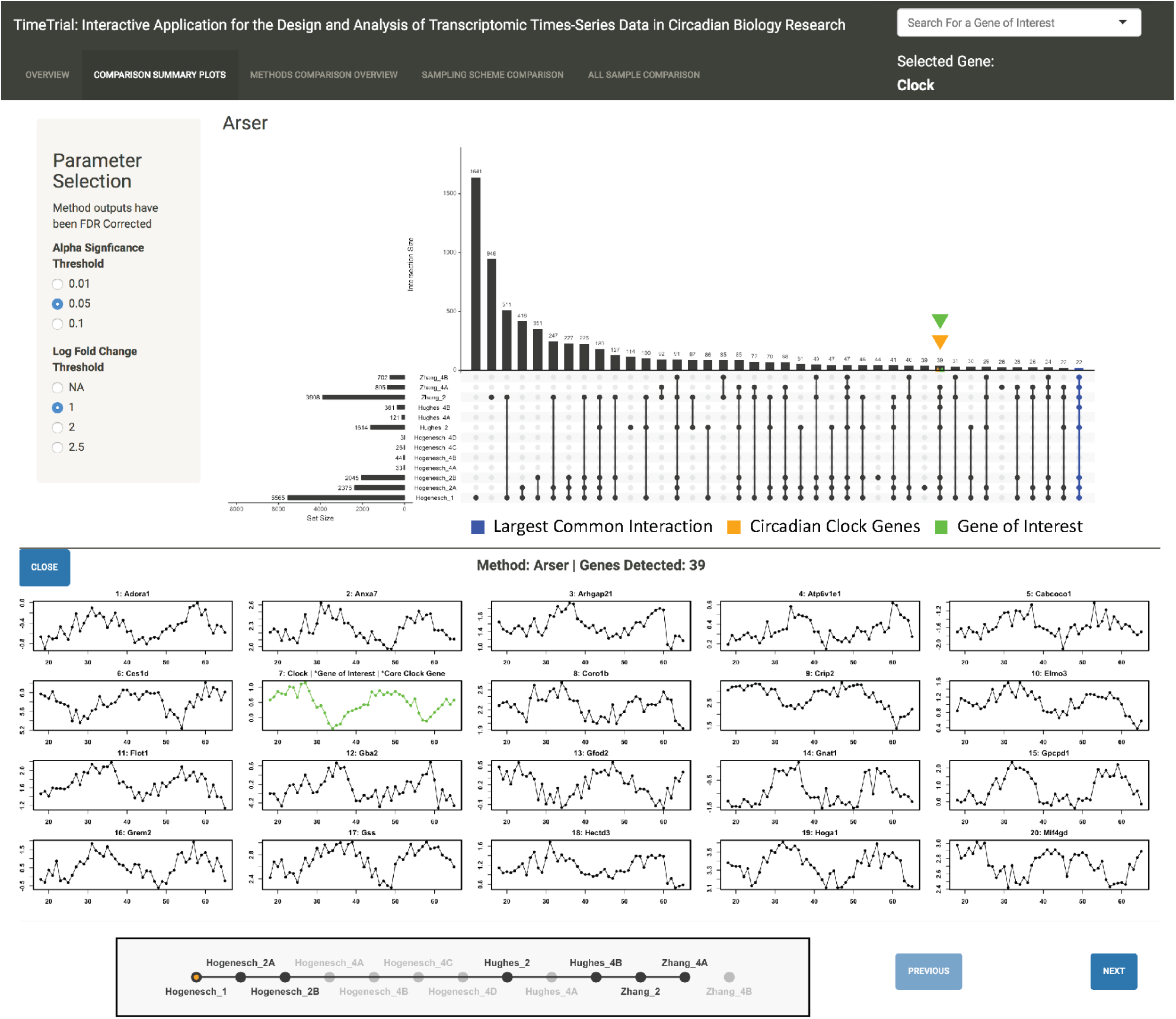
Real Data - TimeTrial: Exploring Processed Data. The real dataset version of TimeTrial allows users to directly compare methods and sampling schemes. Users can set different significance and log fold change thresholds to explore different methods ability to pick up circadian clock genes (orange triangle) and genes of interests (green triangle) across datasets. See (https://github.com/nesscoder/TimeTrial and/or https://nesscoder.shinyapps.io/TimeTrial_Real/) for interactive plots and a complete tutorial.

#### Designing Circadian Experiments with TimeTrial

Most importantly, TimeTrial enables users to develop their own custom sampling scheme for cycle detection (**Figure 6**). While sampling every hour for 48 hours would be “ideal” for cycling detection, it is also expensive. TimeTrial allows the user to explore how scheduling fewer samples will affect the results and explore whether enhanced sampling at specific times of day can improve detection. Irregular sampling schemes may be beneficial for scientific or practical considerations. For instance, one may be interested in monitoring not only cycling genes, but how their dynamics change immediately following an exposure; in this case, one might wish to bias samples toward the time immediately following the stimulus, at the expense of fewer samples later in the time course. The impact of these choices can be explored using TimeTrial’s downsampling tool to test how this alternative sampling scheme compares to the ideal sampling scheme. (NB: The custom analysis is performed using only JTK_Cycle and RAIN, since ARSER and BooteJTK require regularly spaced samples; see **Supplementary Table 1**.) Finally, users can further explore how adjusting the times and spacing of sampling might improve detection and query genes of interest to determine if a specific gene and/or core clock genes are detected.

**Figure 6:**
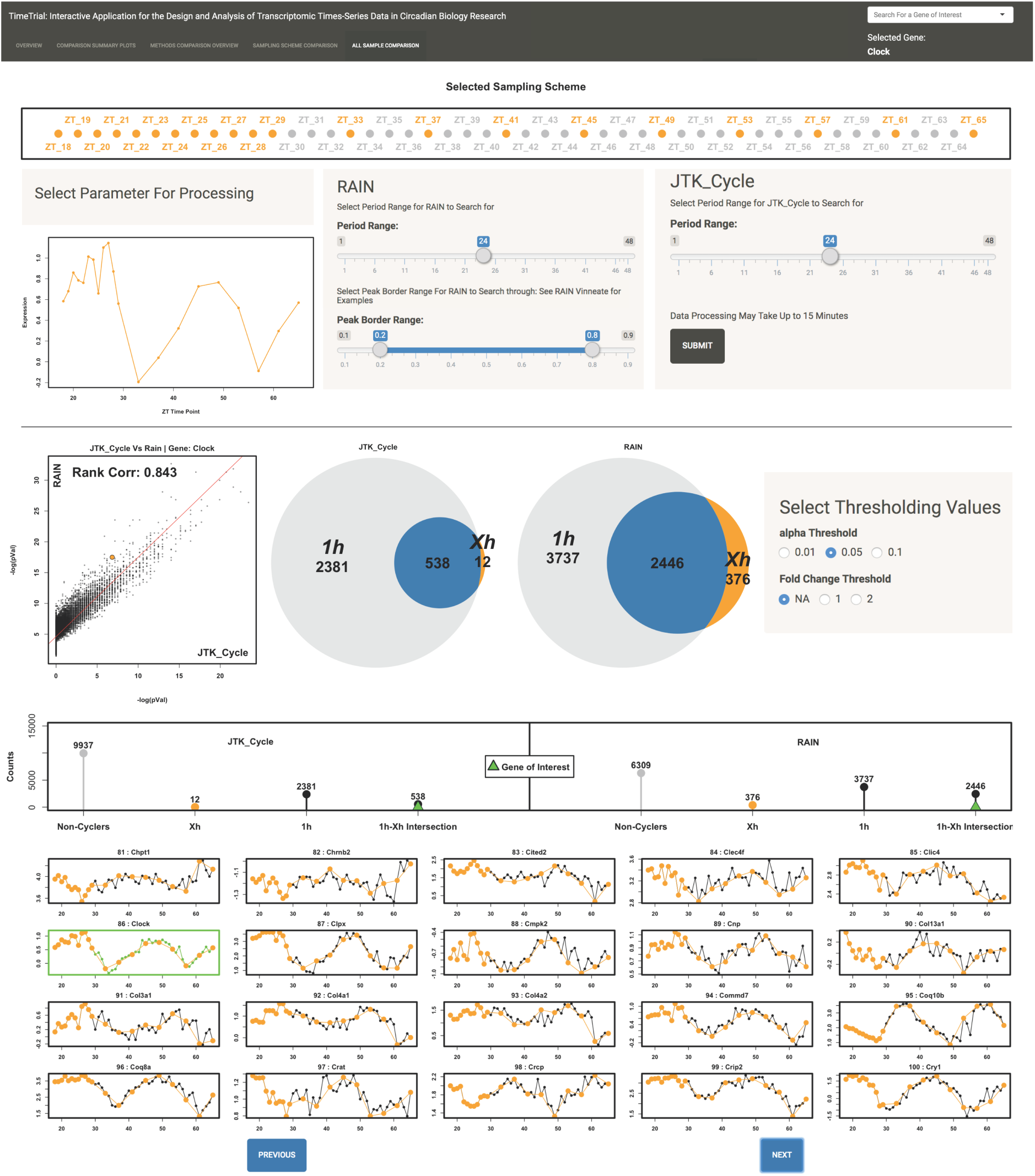
Real Data - TimeTrial: Testing Arbitrary Sampling Schemes. The real dataset version of TimeTrial allows users to define their own custom downsampled sampling scheme and compared the results to that of sampling every 1 hour for 48 hours. The custom sampling scheme analysis is performed using only JTK_Cycle and RAIN, since ARSER and BooteJTK cannot handle uneven sampling (**Supplementary Table 1**). Users can further set different significance and log fold change thresholds and query for genes of interest. See (https://github.com/nesscoder/TimeTrial and/or https://nesscoder.shinyapps.io/TimeTrial_Real/) for interactive plots and a complete tutorial.

The TimeTrial application is freely available for download at https://github.com/nesscoder/TimeTrial, and may also be accessed through a web interface hosted on shinyapps.io: https://nesscoder.shinyapps.io/TimeTrial_Synthetic/, https://nesscoder.shinyapps.io/TimeTrial_Real/. (Note that we recommend local installation from GitHub for greater speed and reliability, as it does not depend on the user’s internet connection.). Full documentation and walk-through tutorials are provided to guide users in the optimal use of these tools.

## Discussion

We developed TimeTrial, a tool that uses synthetic and real data to assist researchers in the design and analysis of circadian time-series experiments that optimize cycling detection. We applied TimeTrial to explore the effects of experimental design on signal shape, examine cycling detection reproducibility across biological datasets, and optimize experimental design for cycle detection. By comparing the performance of different cycling detection algorithms under different sampling schemes, TimeTrial provides valuable guidance for the design of rigorous, reproducible cricadian transcriptomics studies. We expect that these results will be of interest to experimentalists and computational researchers alike.

From an experimental perspective, our results suggest several guidelines for designing circadian time– series studies. First, we demonstrated that 24 hour sampling with concatenation introduces biases which increases the number of false positives, and therefore this practice should be avoided (Hughes et al., 2017). Moreover, our findings suggest that 2-hour resolution is required at a minimum to pick up the dynamical transcriptional changes that occur on the 24 hour circadian scale (corroborating earlier findings (Hughes et al., 2017, 2009)), and that a 2-hour resolution with a single replicate is advantageous over 4-hour resolution in duplicate. We note that the errors produced with the 4-hour schemes include both false negatives (missed cyclers) and false positives (non-cyclers erroneously classified as cycling), compromising the reliability of results obtained from 4-hour sampling schemes. Finally, we observe that different detection methods exhibit different performance depending on the underlying waveform shape, suggesting that the researcher should consider the patterns of interest when selecting an analysis method. For instance, a researcher may decide that classifying a waveform with a strong linear drift as cyclic may be (un)desirable, in which case a method that calls these patterns as (non)cycling should be selected.

From a computational perspective, our results provide a means to benchmark new methods based on their accuracy in synthetic data and reproducibility in real data. They also indicate methodological gaps. Notably, our findings did not define a clear overall “winner” amongst the methods tested, suggesting that there is still a need for methods that perform consistently well in multiple conditions for a variety of waveform shapes. Amongst these findings, we note that no method detects as cycling “contractile” waveforms in which the period changes (as might be the case when an environmental change is introduced), indicating the need for new methods where such patterns are of interest. Our results also highlight the need for methods that have more robust performance with 4–hour sampling designs; the development of such a method would make future experiments more feasible and enable re-analysis of existing data. Finally, we note that any new method should be designed to handle biological and technical replicates in a justifiable way (without requiring concatenation, and ideally without averaging so that the full information about the variance in the data is retained); permit missing data and/or uneven sampling; and be computationally efficient (**Supplementary Table 1**).

Researchers should be aware of read–depth considerations when performing cycling detection using next-generation sequencing. Our analysis was performed on publicly available microarray data, and thus read depth was not considered as a factor in the present analysis. However, previous work has recommended optimal read depths for cycling detection: ∼ 10 million reads per sample to detect >75% of cycling transcripts in fly RNA-seq studies, and ∼ 40 million reads per sample for studying mammals (Li et al., 2015). In the context of TimeTrial, read depth will effect the cost of sampling, and thus acts as a constraint on the number of samples a researcher has at their disposal. Once sample number is determined, TimeTrial can be used to help determine the optimal sampling scheme given this constraint. As more next-generation circadian times-series sequencing data becomes publicly available, future versions of TimeTrial will include the effects of read depth by allowing users to vary this parameter.

Our benchmarking approach is unique in that it provides an assessment of cycling detection for arbitrary sampling schemes. Previous studies were done using fixed sampling frequencies, number of replicates, and sampling lengths. Our analysis used a mixture of different sampling resolutions, number of replicates, sampling lengths, and noise levels. Ultimately, we attempted to model biological experimental practice, where a fixed sampling scheme is not always possible as a result of monetary constraints. Our findings suggest that the sampling schemes and signal shape, rather than the cycling detection method, will have the largest impact on cycle detection. Thus, in designing a time-series experiment, researchers should contemplate the total number of samples at their disposal; how those samples should be used across replicates, length, and intervals; and how the chosen scheme allows cycling detection methods to pick up different signal shapes (i.e. symmetric, non-symmetric, peak, trends, etc.).

Additionally, our benchmarking approach is unique in that it explicitly considers the reproducibility of cycling detection results. By considering the concordance of genes detected as cycling across multiple independent datasets, we directly assess whether genes detected in one study would be validated in another. We propose that this assessment of reproducibility, rather than the number of cycling genes detected, should be the standard against which new methods are judged.

## Methods

### Generating Synthetic Datasets

240 unique synthetic time course datasets each comprising 11,000 expression profiles were generated in R. Each dataset consisted of different number of replicates (1, 2, 3), sampling intervals (2h, 4h, 6h, 8h), sampling lengths (24h, 48h, 72h, 96h), and noise levels as a percent of the wave form amplitude (0%, 10%, 20%, 30%, 40%) (**Figure 1A**).

Within each condition, eleven base waveforms were simulated to mimic expression patterns observed in nature: periodic patterns, non-periodic patterns, and dynamics that have a cyclic component but do not meet the strict definition of periodicity. Seven of these eleven shapes were considered **Cyclic** (Sine, Peak, Sawtooth, Linear Trend, Damped, Amplified, Contractile); four were considered **Non-Cyclic** (Flat, Linear, Sigmoid, and Exponential); examples are given in **Figure 1B**. For each waveform in each condition, 1000 “genes” were simulated with varying amplitudes, phases, and shape parameters (e.g., the envelope for damped/amplified waves), yielding in total 11,000 simulated genes for each of the 240 conditions. Variation in amplitude of underlying functions where drawn from a log normal uniform distribution with mean 1.302 and standard deviation 0.30, as modeled from real data amplitude distributions to simulate differences in amplitude between genes. Additional variation in phase of underlying functions where drawn from a uniform distribution between 0 and 2*π*, to simulate differences in phase between genes. The data was mean centered, as is common in pre-processing for cycle detection. A complete list of the waveform function definitions and source code for generating the data can be found in the *supplement*.

### Pre-Processing Microarray Datasets

CEL files from three mouse liver Affymetrix microarray time-series expression sets (Hogenesch 2009 - GSE11923 (Hughes et al., 2009), Hughes 2012 - GSE30411 (Hughes et al., 2012), Zhang 2014 - GSE54650 (Zhang et al., 2014)) were downloaded from the Gene Expression Omnibus database (GEO). In each experiment, wildtype C57BL/6J mice were entrained to a 12 h light, 12 h dark environment before being released into constant darkness. Mouse age, length of entrainment, time of sampling, and sampling resolution vary by experiment. The data was subsequently normalized by Robust Multi-array Average (rma) using the Affy R Package (Gautier et al., 2004) and checked for quality control using the Oligo R Package (Carvalho and Irizarry, 2010), following each package’s vignette respectively. Since each GEO dataset used a different microarray platform (affy_mouse430_2, affy_moex_1_0_st_v1, affy_mogene_1_0_st_v1), each had a different set of probes. A common set of features needed to be identified in order to compare across microarrays. Probes for each dataset where mapped to genes based on pre-aligned databases specific to each microarray (mouse4302.db, moex10sttranscriptcluster.db, mogene10sttranscriptcluster.db). Multiple probes corresponding to one gene were aggregated by taking the mean expression. A final 12868 common set of genes across all three microarray platforms were used for subsequent analysis. See *supplement* for code.

### Processing Microarray Datasets

To characterize the effects of sampling schemes using real data, the three datasets were downsampled to simulate the effects of sampling at 2-hours and 4-hour intervals. The Hughes 2012 and Zhang 2014 datasets were sampled every 2h for 48 hours. Each of these experimental time-series were downsampled to every 4 hours to generate 4 additional time-series datasets. The Hogenesch 2009 dataset, sampled every hour for 48 hours, was downsampled to 2 datasets every 2 hours and 4 datasets sampled every 4 hours to generate 6 additional time-series datasets. Ultimately, 13 datasets (3 original and 10 downsampled) were processed by all four cycling detection methods (ARSER, BooteJTK, JTK_Cycle, and Rain), using each method’s recommended parameter settings as defined by the sampling length and interval (**Figure 1C**). We thus aimed to assess a method’s robustness by the ability to consistently score genes as cycling versus non-cycling across experimental datasets and down-samplings. A complete list of the experimental parameters and source code can be found in the *supplement*.

### Application of Cycling Detection Algorithms

All datasets were processed by all four cycling detection methods (JTK_Cycle (Hughes et al., 2010), ARSER (Yang and Su, 2010), RAIN (Thaben and Westermark, 2014), and BooteJTK (Hutchison et al., 2018); **Supplementary Table 1**), using each method’s recommended parameter settings as defined by the sampling length and interval. Since ARSER and BooteJTK do not have a built in function for dealing with replicates, replicates were either averaged together or concatenated, following the two common practices in the field. JTK_Cycle and Rain used the replicate procedures recommended in their documentation. A complete list of the experimental parameters and source code can be found in the *supplement*.

## Acknowledgements

Research reported in this publication was supported by the NSF-Simons Center for Quantitative Biology at Northwestern University, an NSF-Simons MathBioSys Research Center. This work was supported by a grant from the Simons Foundation/SFARI (597491-RWC) and the National Science Foundation (1764421). The content is solely the responsibility of the authors and does not necessarily represent the official views of the National Science Foundation and Simons Foundation.

## Notes

Author contributions: E.N.C. and R.B. designed research; E.N.C., M.I., W.L.K., R.A., and R.B. contributed to the design requirements of TimeTrial; E.N.C developed the TimeTrial software, analyzed the data, and produced the tutorials; E.N.C and R.B. wrote the paper; all authors read and approved the final manuscript.

