## Supplementary for "TimeTrial: An Interactive Application for Optimizing the Design and Analysis of Transcriptomic Times-Series Data in Circadian Biology Research"

### JTK_Cycle

JTK_Cycle [1] is a non-parametric method based on the Jonchkeere-Terpstra and Kendall rank based sum test. Implemented in the R programming language, JTK_Cycle uses the Jonchkeere-Terpstra test to detect the monotonic orderings of data across ordered independent groups and subsequently uses the Kendall rank sum test to compare the experimental Jonchkeere-Terpstra statistic to that of a reference curve. JTK_Cycle strength is the method has built in structures to handle uneven sampling, missing data, and replicate samples (**Table 1**). In practice, all of these feature are important to researchers. Often data points are missed as a result of sequencing errors [2]. Moreover, uneven sampling and replicates are useful in the design of time course experiment where researchers may want dense sampling at a specific stretch of the time course to detect a signal of interest and subsequent sparser sampling as the time course progresses. While JTK_Cycle thus allows for flexibility in experimental design, results show that by using a reference waveform (i.e. a cosine wave), JTK_Cycle is biased towards that reference and periodic signals that do not fit the reference pattern (i.e. a sawtooth wave) can go undetected [3].

### ARSER

ARSER [4] is a parametric method based on autoregressive spectral estimation. Implemented in both Python and the R programming language, by first detrending and then smoothing the data, ARSER assesses statistical significance of the waveforms fit to a sinusoidal curve. As a nature of the detrending in the algorithm, Arser has been shown to pick up trending oscillatory dynamics such as linear trends [4]. ARSER however does not have built in structures to handle missing data, uneven sampling, or replicate samples (**Table 1**).

### RAIN

RAIN [3] is a non-parametric method based on the rank test for umbrella alternative – a generalization of the Jonckheere-Terpstra test – that does not require a user defined reference signal. Implemented in the R programming language, RAIN was developed to expand the range of signals detected by JTK_Cycle. Using the rank test for umbrella alternative, RAIN searches for monotonic patterns of rising followed by monotonic patterns of falling, but does not assume any relationship between the patterns. As such, a user-defined reference signal is not employed, allowing for the detection of rhythms that other referenced based methods may miss. RAIN further has built in structures to handle uneven sampling, missing data, and replicate samples (**Table 1**).

### BooteJTK

BooteJTK [5] is a non-parametric method based on an empirical Bayes procedure. Implemented in Python programming language, BooteJTK "shrinks" the spread in variances across time points, generates bootstrap time-series from the variance estimations, and computes statistical significance of cycling by averaging the results of the bootstrapped time-series via a non-parametric pairwise rank order correlation, Kendall's $\tau$, in relation to a gamma distribution. BooteJTK uses a range of phase shifted reference waveforms to avoid bias toward a single reference signal. Furthermore, the variance shrinking coupled with the bootstrapping procedure is argued to improve consistency across related experimental datasets [5]. Nonetheless, the method can be computationally expensive. As dataset size scales, the bootstrapping procedure become computationally more costly. Furthermore, the method does not have built in structures to handle uneven sampling or deal with replicates. The method can handle missing data, as long as there is at least some data present for every time point sampled. Since BooteJTK groups ZT time points across period cycles (i.e. ZT_2 groups with ZT_26), if the grouped ZT times are missing across period cycles – more characteristic of uneven sampling – BooteJTK's detection procedures breaks down (**Table 1**).

| Method | Replicates | Missing Data | Uneven Sampling | Efficiency | Programming Language |
|---|---|---|---|---|---|
| ARSER | - | - | - | ✓ | Python/R |
| BooteJTK | - | ✓ | - | - | Python |
| JTK_Cycle | ✓ | ✓ | ✓ | ✓ | R |
| RAIN | ✓ | ✓ | ✓ | ✓ | R |

**Table 1: Comparison of Method Features**
(✓) represent the method is implementation to handle the feature, while (-) denotes a method does not have a valid implementation. Missing data refers to sporadic non-sequenced timepoints on a per gene basis, while uneven sampling refers to a systematic omission of a specific timepoint across all genes in the time-series. Thus, BooteJTK has a (✓) for missing data, since the imputation procedure can handle the sporadic level of missingness; but a (-) for uneven sampling, since the imputation procedure fails with the systematic omission of a particular time point across all genes.

### Parameters Synthetic Data | methodParameters_SyntheticData.xlsx

The excel file contains all the parameters used for processing each synthetic datasets. The spreadsheet is broken down by method. All datasets were processed by all four cycling detection methods (Arser, BooteJTK, JTK_Cycle, and Rain), using each method's recommended parameter settings as defined by the sampling length and interval. Since Arser and BooteJTK do not have built in function for dealing with replicates, replicates where either averaged together or concatenated as are the two common practices in the field. JTK_Cycle and Rain used the replicate procedures recommended in their documentation. See (`https://github.com/nesscoder/TimeTrial`) for source code and additional files.

### Parameters Biological Data | methodParameters_BiologicalData.xlsx

The excel file contains all the parameters used for processing each synthetic datasets. The spreadsheet is broken down by method. All datasets were processed by all four cycling detection methods (Arser, BooteJTK, JTK_Cycle, and Rain), using each method's recommended parameter settings as defined by the sampling length and interval. See (`https://github.com/nesscoder/TimeTrial`) for source code and additional files.

### Synthetic Data | Function Form

| Cycling Waveform | Signal Function |
|---|---|
| Sine | $f(t) = \frac{A}{2}\sin\left(\frac{2\pi}{24}(t-\phi)\right) + \epsilon$ |
| Peak | $f(t) = A\left(\left\|\sin\left(\frac{2\pi}{24}(t-\phi)\right)\right\|^p\right) + \epsilon\,;\ p \in [10, 60]$ |
| Sawtooth | $f(t) = A\min\left(\frac{(t \mod 24)}{\phi}, \frac{(t \mod 24) - 24}{\phi - 24}\right) + \epsilon$ |
| Linear Trend | $f(t) = lt + \frac{A}{2}\sin\left(\frac{2\pi}{24}(t-\phi)\right) + \epsilon\,;\ l \in [-2, 2]$ |
| Damped | $f(t) = e^{-dt} \cdot \frac{A}{2}\sin\left(\frac{2\pi}{24}(t-\phi)\right) + \epsilon\,;\ d \in [0.01, 0.03]$ |
| Amplified | $f(t) = e^{at} \cdot \frac{A}{2}\sin\left(\frac{2\pi}{24}t-\phi\right) + \epsilon\,;\ a \in [0.01, 0.015]$ |
| Contractile | $f(t) = \frac{A}{2}\sin\left(\frac{2\pi}{24}\left(t^k/24-\phi\right)\right) + \epsilon\,;\ k \in [1.8, 1.9]$ |

**Table 2: Cycling Waveform Functions**
The time, $t$, is evenly distributed every 2 hours between $t = 0$ and $t = 96$. Phases $\phi$ are measured in hours, and were uniformly sampled. All waveforms were down-sampled to account for the different sampling lengths (24, 48, 72, 96) and sampling intervals (2, 4, 6, 8). The amplitudes, $A$, was drawn from a log-normal distribution with $\mu = 1.302$ and $\sigma = 0.303$, as derived from the distribution of amplitudes in observed biological data. Gaussian white noise $\epsilon$ was added to each point independently as a percentage of the amplitude, $\epsilon = rA\eta$ where $\eta \sim \mathcal{N}(0, 1)$ and $r = \{0\%, 10\%, 20\%, 30\%, 40\%\}$. All other parameters were uniformly sampled over the ranges denoted in the table. Range bounds were chosen to maintain the mean expression and standard deviation seen in real data. See (`https://github.com/nesscoder/TimeTrial`) for source code and additional files.

| Non-Cycling Waveform | Signal Function |
|---|---|
| Flat | $f(t) = A + \epsilon$ |
| Linear | $f(t) = mt + \epsilon\,;\ m \in [-5, 5]$ |
| Sigmoid | $f(t) = \frac{A}{1 + e^{g(t-\phi)}} + \epsilon\,;\ g \in [-1, 1]$ |
| Exponential | $f(t) = \frac{A}{100}e^{dt} + \epsilon\,;\ d \in [0.09, 0.05]$ |

**Table 3: Non-Cycling Waveform Functions**
The time, $t$, is evenly distributed every 2 hours between $t = 0$ and $t = 96$. All waveforms were down-sampled to account for the different sampling lengths (24, 48, 72, 96) and sampling intervals (2, 4, 6, 8). The amplitudes, $A$, was drawn from a log-normal distribution with $\mu = 1.302$ and $\sigma = 0.303$, as derived from the distribution of amplitudes in observed biological data. Gaussian white noise $\epsilon$ was added to each point independently as a percentage of the amplitude, $\epsilon = rA\eta$ where $\eta \sim \mathcal{N}(0, 1)$ and $r = \{0\%, 10\%, 20\%, 30\%, 40\%\}$. All other parameters were uniformly sampled over the ranges denoted in the table. Range bounds were chosen to maintain the mean expression and standard deviation seen in real data. See (`https://github.com/nesscoder/TimeTrial`) for source code and additional files.

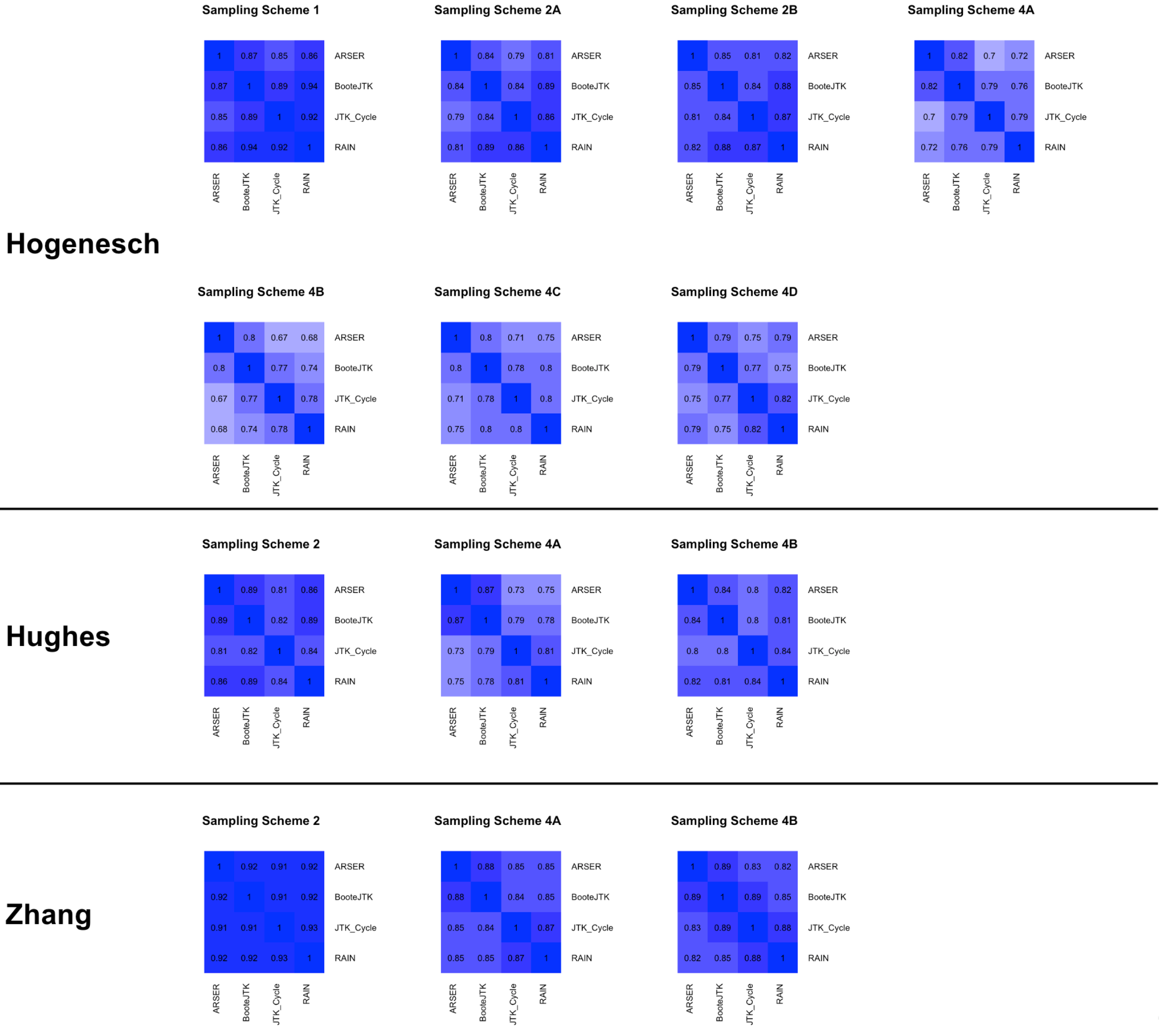


**Figure 1: Method Comparison** | The spearman rank correlation, ($\rho$), of the resultant raw p-values in each dataset with a given sampling scheme were computed. This correlation compares the ranking of genes from one method to their corresponding ranking in a different method. See (`https://github.com/nesscoder/TimeTrial`) for interactive plots and a complete tutorial.

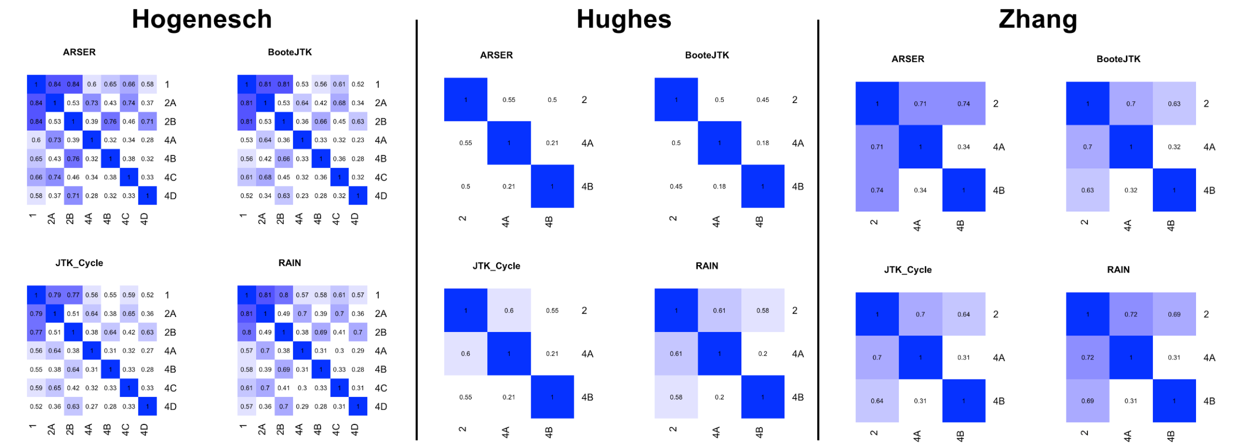


**Figure 2: Sampling Schemes Comparison** | The spearman rank correlation, ($\rho$), of the resultant raw p-values in each dataset with a given method were computed. This correlation compares the ranking of genes from one sampling scheme to their corresponding ranking in a different sampling scheme. See (`https://github.com/nesscoder/TimeTrial`) for interactive plots and a complete tutorial.
